# Transcriptional responses to *in vitro* macrocyclic lactone exposure in *Toxocara canis* larvae using RNA-seq

**DOI:** 10.1101/2024.12.20.629602

**Authors:** Theresa A. Quintana, Matthew T. Brewer, Jeba R. Jesudoss Chelladurai

## Abstract

*Toxocara canis*, the causative agent of zoonotic toxocariasis in humans, is a parasitic roundworm of canids with a complex lifecycle. While macrocyclic lactones (MLs) are successful at treating adult *T. canis* infections when used at FDA-approved doses in dogs, they fail to kill somatic third-stage larvae. In this study, we profiled the transcriptome of third-stage larvae derived from larvated eggs and treated *in vitro* with 10 µM of the MLs – ivermectin and moxidectin with Illumina sequencing. We analyzed transcriptional changes in comparison with untreated control larvae. In ivermectin-treated larvae, we identified 608 differentially expressed genes (DEGs), of which 453 were upregulated and 155 were downregulated. In moxidectin-treated larvae, we identified 1,413 DEGs, of which 902 were upregulated and 511 were downregulated. Notably, many DEGs were involved in critical biological processes and pathways including transcriptional regulation, energy metabolism, neuronal structure and function, physiological processes such as reproduction, excretory/secretory molecule production, host-parasite response mechanisms, and parasite elimination. We also assessed the expression of known ML targets and transporters, including glutamate-gated chloride channels (GluCls), and ATP-binding cassette (ABC) transporters, subfamily B, with a particular focus on P-glycoproteins (P-gps). We present gene names for previously uncharacterized *T. canis* GluCl genes using phylogenetic analysis of nematode orthologs to provide uniform gene nomenclature. Our study revealed that the expression of *Tca-glc-3* and six ABCB genes, particularly four P-gps, were significantly altered in response to ML treatment. Compared to controls, *Tca-glc-3*, *Tca-Pgp-11.2*, and *Tca-Pgp-13.2* were downregulated in ivermectin-treated larvae, while *Tca-abcb1*, *Tca-abcb7*, *Tca-Pgp-11.2*, and *Tca-Pgp-13.2* were downregulated in moxidectin-treated larvae. Conversely, *Tca-abcb9.1* and *Tca-Pgp-11.3* were upregulated in moxidectin-treated larvae. These findings suggest that MLs broadly impact transcriptional regulation in *T. canis* larvae.

## Introduction

*Toxocara canis* is a parasitic roundworm that infects canids. Zoonotic *T. canis* larval infection in humans, known as toxocariasis, is a significant public health concern [1]. In the U.S., it is estimated that 5% to 20% of the population (∼17 – 66 million people) have been exposed to *Toxocara* spp., with notably higher exposure among individuals from lower socioeconomic backgrounds and those residing in the South and Northeastern parts of the country [2, 3]. On a global scale, the estimated number of people exposed is 19% (1.52 billion people)[4]. The risk of infection with *T. canis* is especially high in developing countries where poor sanitation and animal contact are common [5].

Humans become infected by accidentally consuming larvated eggs of *T. canis* [6]. Toxocariasis in humans has various clinical manifestations, ranging from infections with mild symptoms such as fever and abdominal pain to severe visceral and/or ocular larva migrans [7]. To effectively mitigate the impact of toxocariasis on human health [8], it is crucial to gain a deeper understanding of the biological processes and pathways involved in the transmission and infection of *T. canis*.

The lifecycle of *T. canis* involves the development of adult worms in the small intestine of infected dogs. Infection in dogs typically begins by the ingestion of larvated *T. canis* eggs found in contaminated soil or on the fur of infected animals [5]. Ingested eggs hatch in the duodenum, penetrate the intestinal mucosa, and migrate to the liver and lungs [9]. In dogs under three months of age, larvae are coughed up from the lungs and swallowed, completing their lifecycle to adulthood in the small intestines. Adult female worms shed thousands of eggs, which pass out in the feces and contaminate the surrounding environment. These eggs can survive in the environment for long periods of time, creating an increased exposure risk for humans [10].

Impacts of *T. canis* infection in dogs can vary depending on the severity of the infection and the age of the animal [11–14]. In severe infections, adult *T. canis* infection can lead to weight loss, poor growth, vomiting, and diarrhea [9]. Adult *T. canis* infections can be successfully treated using several anthelmintic drug classes including tetrahydropyrimidines (pyrantel pamoate, 5 mg/kg [15]), benzimidazoles (fenbendazole, 50 mg/kg [16], or febantel, 25 mg/kg [17]) and macrocyclic lactones (MLs), such as ivermectin (200 micrograms/kg [18]), moxidectin (2.5 mg/kg [19]), and milbemycin oxime (0.5 mg/kg [20]). In older dogs, larvae undergo somatic migration, commonly arresting in the skeletal muscle, kidneys, and liver [9, 21]. Somatic larvae can remain dormant for long periods and reactivate in dams during the third trimester of pregnancy [22–25]. Larvae are transmitted *in utero* from the infected dam to developing neonates [26]. Thus, somatic larvae in a dog’s muscles and viscera remain a persistent source of infection. Arrested somatic larvae of *T. canis* in dogs cannot be killed by anthelmintic treatments, including MLs, when used at FDA-approved labeled doses and therefore remain arrested in the tissues, demonstrating a perceived tolerance to the therapeutic effects of these compounds [27].

The mechanism behind the tolerance of *T. canis* somatic larvae to MLs is not fully understood. MLs act on glutamate-gated chloride channels (GluCls) in parasitic nematodes [28, 29]. GluCls are part of the cys-loop ligand-gated ion channels (LGICs) superfamily and play an important role in signal transduction in the invertebrate nervous system [30]. GluCls consist of five heterogeneous or homogeneous subunits [31]. Downregulation in the transcription of GluCls is associated with ivermectin resistance in the parasitic nematode *Haemonchus contortus* [32, 33]. It is thought that alterations in GluCl subunit expression lead to different ion channel stoichiometry thereby altering ion channel pharmacology [34]. However, neither this phenomenon nor transcriptional changes in *T. canis* GluCls in response to ML treatment have been studied.

Besides alterations in the GluCl target sites, there are other non-specific mechanisms that may protect nematodes from MLs. In several nematodes, Permeability glycoproteins (P-gps), which belong to the ATP-binding cassette (ABC) transporter subfamily B (ABCB1) are involved with ML efflux, which is responsible for phenotypic resistance [34–44]. In *T. canis*, *Tca-Pgp-11* has been demonstrated to transport ivermectin and be inhibited by known P-gp inhibitors [45]. Additionally, thirteen P-gp genes are expressed in various larval stages of *T. canis* and have been studied using qPCR [46].

To better understand the mechanisms underlying the effect of MLs on *T. canis*, a comprehensive analysis of its transcriptome is essential. Currently, there is limited data on the transcriptomes of *T. canis*, with one previous study comparing transcription in adult and larval isolates [47] and another comparing transcription in adult male and female isolates [48]. Since the infectious stage of *T. canis* is the third-stage larvae (L3), there is a critical need to understand the transcriptional effects of MLs on this stage. Additionally, L3 larvae from eggs serve as a surrogate for somatic L3s, which are difficult to isolate from the tissues of infected dogs.

In this study, we assessed the differential expression of genes in the comprehensive transcriptome of hatched *T. canis* third-stage larvae treated with the MLs ivermectin or moxidectin. We performed gene ontology analyses on the differentially expressed genes (DEGs). Additionally, we analyzed the transcription of annotated GluCl and ABCB genes of *T. canis* in response to ivermectin and moxidectin.

## Materials and Methods

### Parasites

Adult *T. canis* male and female worms were opportunistically obtained from a naturally infected dog that was presented to the Kansas State University (KSU) Veterinary Diagnostic Laboratory for necropsy. Eggs were isolated from the uterus of adult gravid female worms and incubated at 25°C for three weeks to allow larval development to L3s. Hatching was induced as previously described [49], larvae were isolated and suspended in Roswell Park Memorial Institute (RPMI) –1640 medium (Gibco by Thermo Fisher Scientific, Grand Island, NY). Hatched larvae (n = ∼500 – 1,000) were subjected to three conditions: incubation with 10 µM ivermectin (IVM, Thermo Fisher Scientific, Ward Hill, MA), 10 µM moxidectin (MOX, Sigma-Aldrich, St. Louis, MO), or no treatment (controls, RPMI-1640 only), at 37°C for 12 hours and in biological triplicates. There was no observed loss in larval motility following drug incubation compared to controls. Samples were frozen at –80°C until RNA extraction.

### RNA isolation and RNA-seq

Larvae were homogenized and RNA was isolated from each sample using the standard Trizol reagent extraction protocol with Phasemaker tubes (Invitrogen, Carlsbad, CA) as recommended by the manufacturer. Total RNA was quantified using a Qubit RNA Broad Range Assay Kit (Life Technologies/ Invitrogen, Eugene, OR). RNA-seq was performed at the CEZID Molecular and Cellular Biology Core at KSU. Briefly, stranded mRNA was purified with oligo(dT) magnetic beads and libraries were prepared with an Illumina® Stranded mRNA Prep, Ligation kit (Illumina, San Diego, CA) with a minimum of 100 ng of RNA. A brief graphical representation of these methods can be found in Figure 1. Paired-end sequencing (2×75 cycles) was performed on an Illumina NextSeq 550 sequencer using a NextSeq 500/550 High Output Kit v2.5 (Illumina, San Diego, CA).

**Figure 1.**
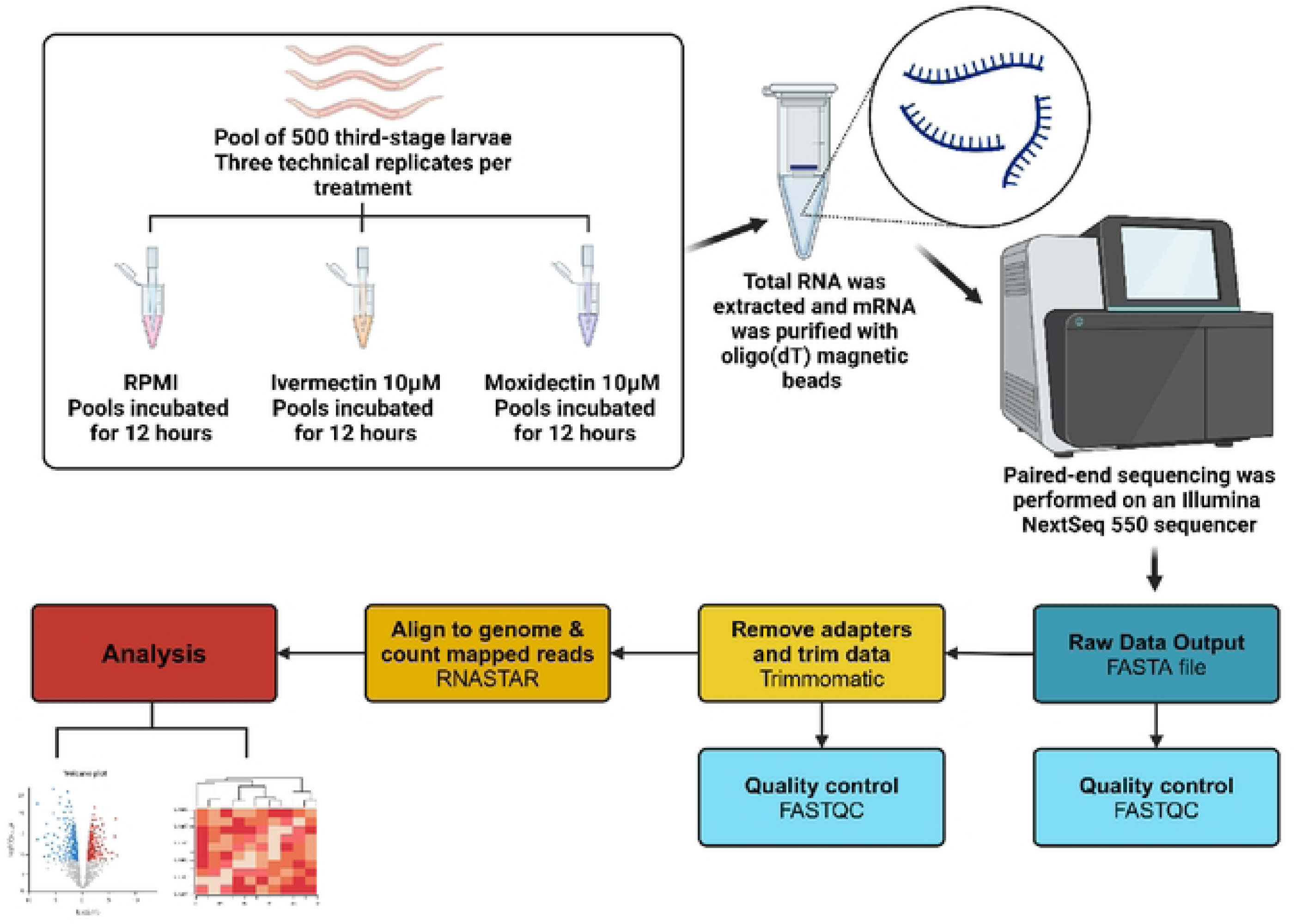
Brief graphical depiction of the experimental methods for the transcriptomic study of *Toxocara canis* third-stage larvae exposed to ivennectin, moxidectin, or RPMl-1640 (controls). ll1is figure was created using Biorender.com.

### Bioinformatic Analysis

Raw data was obtained in FASTQ format and analyzed using the STAR pipeline on Galaxy servers [50]. Briefly, FASTQ files were trimmed using Trimmomatic (Version 0.38) [51]. Trimmed reads were aligned and mapped to the genes annotated in the *T. canis* draft genome (NCBI GenBank assembly GCA_000803305.1), and gene expression was measured as reads per gene using STAR [52]. Raw and trimmed read qualities were assessed with FastQC (Version 0.12.1). Alignment quality was assessed with Qualimap BAMQC (Version 2.2.2c) [53, 54]. Reads per gene counts from STAR were analyzed using custom R scripts (Version 4.3.2). Gene expression normalization, differential expression analysis in response to drug treatment, and principal components analysis (PCA) were performed using *DESeq2* (Version 1.40.1). Volcano plots were created with *EnhancedVolcano* (Version 1.18.0). Pearson correlations as a measure of intragroup invariance were calculated and differentially expressed genes were filtered with *tidyverse* (Version 2.0.0). Venn diagrams to determine differentially expressed genes that were shared between drug treatments were created with *ggvenn* (Version 0.1.10). Functional gene ontology analysis was plotted with *gprofiler2* (Version 0.2.2) [55]. Gene set enrichment on differentially expressed genes was performed in ShinyGO (Version 0.80) [56].

Gene-specific information about 27 ML-associated genes (transporters and targets) was obtained using *rentrez* (Version 1.2.3) [57]. Phylogenetic analysis was performed using NGPhylogeny.fr [58, 59]. Briefly, protein sequences of GluCls were manually curated from the Genbank protein database, aligned with MAFFT [60], and trimmed with BMGE [61]. A maximum likelihood (ML) tree was constructed with PhyML 3.0 [62] after determining the best model using SMS [63]. The tree in Newick format was visualized using iTOL (version 6) [64, 65] and *ggtree* (Version 3.10.1) [66]. Heatmaps were created with variance stabilized and transformed DESeq2 data using *ComplexHeatmap* (Version 2.16.0), *ggpubr* (Version 0.6.0), and *grid* (Version 4.3.2). Expression levels of the 27 ML-associated genes were tested for normality with the Shapiro-Wilk test using JASP (Version 0.13.1.0) [67]. Boxplots to visualize expression levels were created using *ggplot2* (Version 3.5.0), and expressions were compared with the Tukey HSD test using *ggpubr* (Version 0.6.0). Other R libraries used included *ggprism* (Version 1.0.5) and *patchwork* (Version 1.2.0).

## Results

### Sequencing

We obtained transcriptomic sequences from hatched L3s that were untreated, or treated with ivermectin, or treated with moxidectin (n=3, ∼500 larvae each) using an Illumina NextSeq 550 sequencer. A total of 295,503,166 paired-end raw reads were generated, with an average of 32,833,685 reads per sample. After adapter trimming, a total of 293,127,407 pair-end reads were retained, with an average of 32,569,712 reads per sample. The reads were assembled against the annotated reference genome (GenBank Accession #PRJNA248777; 18,596 genes [47]) before transcripts were quantified using STAR. The sequencing data generated was of high quality, with an average of 81.3% of the reads mapping to the *T. canis* draft genome (Table 1).

**Table 1.** Summary of the features of the sequenced cDNA libraries for samples from this study.

| Sample name | Sample group | Paired read count (raw) | Nucleotide count (raw | Paired read count (trimmed) | Nucleotide count (trimmed) | Mapped read count (%) |
| --- | --- | --- | --- | --- | --- | --- |
| Control T1 | Control *T. canis* L3 | 32,249,245 | 2.3 Gbp | 32,174,441 | 2.2 Gbp | 84.54% |
| Control T7 | Control *T. canis* L3 | 54,071,313 | 3.9 Gbp | 54,010,559 | 3.8 Gbp | 86.05% |
| Control T16 | Control *T. canis* L3 | 40,128,609 | 2.9 Gbp | 40,087,855 | 2.8 Gbp | 87.9% |
| Ivermectin T3 | Ivermectin *T. canis* L3 | 22,121,380 | 1.6 Gbp | 21,543,247 | 1.4 Gbp | 81.79% |
| Ivermectin T4 | Ivermectin *T. canis* L3 | 17,179,802 | 1.2 Gbp | 17,105,795 | 1.2 Gbp | 92.92% |
| Ivermectin T16 | Ivermectin *T. canis* L3 | 26,457,311 | 1.9 Gbp | 25,746,262 | 1.8 Gbp | 91.54% |
| Moxidectin T2 | Moxidectin *T. canis* L3 | 29,364,220 | 2.1 Gbp | 29,186,760 | 2.0 Gbp | 92.84% |
| Moxidectin T7 | Moxidectin *T. canis* L3 | 36,440,579 | 2.6 Gbp | 36,389,622 | 2.5 Gbp | 45.54% |
| Moxidectin T8 | Moxidectin *T. canis* L3 | 37,490,707 | 2.7 Gbp | 36,882,866 | 2.6 Gbp | 68.59% |

To determine similarities between the untreated and treated larval groups a principal component analysis (PCA) was conducted to visualize the relationship between the groups based on variance stabilized transform of normalized expression values (Figure 2A). Principal component 1 explained 39% of the variance observed. Principal component 2 explained 29% of the variance observed. Intragroup Pearson correlation within the group ranged from 0.94 to 0.99 for the control group, 0.71 to 0.95 for the ivermectin-treated group, and 0.97 to 0.98 for the moxidectin-treated group. Untreated controls clustered separately from the ivermectin– and moxidectin-treated larvae, indicating that the drug treatments had a significant impact on the gene expression profiles of the larvae.

**Figure 2.**
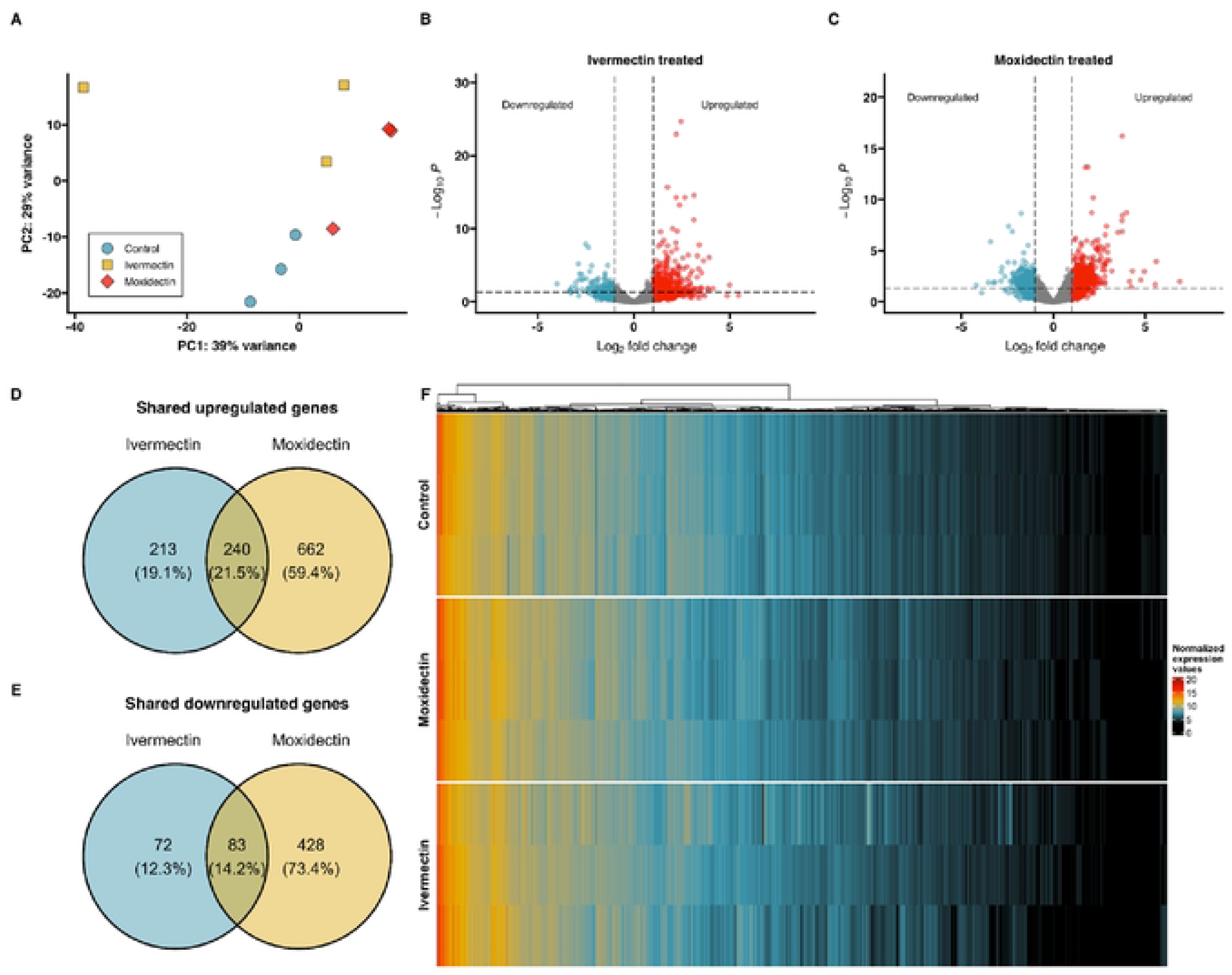
(**A**) Principal Component Analysis (PCA) of *Toxocara canis* third-stage larvae (L3s) exposed to RPMI-1640 medium (controls; blue circle), ivermectin (yellow square),ormoxidectin (red diamond). Each point represents a pool of ∼500 L3s (one biological replicate); **(B-C)** Volcano plots of differentially expressed genes in **(B)** ivcrmcctin-or **(C)** moxidcctin-trcatcd larvae compared to untreated controls. Significance thresholds were set at log2FC > I (two-fold change) for upregulated genes, and log2FC < –I for do,vnregulated genes and adjusted p-value < 0.05. Upregulated genes are shown in red, downregulated genes in blue, and non-significantgenes in gray. A total of 18,596 genes were analyzed, with 454 significantly uprcgulated and 155 significantly downrcgulatcd in ivcrmcctin-treatcd L3s and 902 significantly uprcgulated and 511 significantly downrcgulatcd in moxidcctin-trcated L3s; **(D-E)** Venn diagrams dipicting the overlap of shared upregulated **(D)** and dov.’llregulated **(E)** genes in larvae treated, vith ivermectin and moxidectin. The upregulated genes show an overlap of 240 shared genes, while the do,vnregulated genes sho,v an overlap of83 shared genes; **(F)** Heatmap of normalized expression (vst) values for all 18,596 genes annotated in the *1’. canis* genome for control, ivcrmcctin-, and moxidcctin­ trcated L3s. Color intensity represents normalized expression values, vith the scale ranging from black (lowest expression) to red (highest expression).

### Analysis of Differentially Expressed Transcripts

To identify differentially expressed transcripts, we used the reference-guided transcript assembly method using STAR and DESeq2. Volcano plots showing fold changes and adjusted p-values for the 18,596 genes are shown in Figure 2B-C. We identified 608 differentially expressed genes in larvae treated with ivermectin compared to controls of which 453 were upregulated (two-fold change, that is, log2fold change > 1 and adjusted p-value < 0.05), and 155 were downregulated (log2fold change < –1 and adjusted p-value < 0.05), respectively (Figure 2B). Additionally, we identified 1,413 differentially expressed genes in larvae treated with moxidectin compared to controls, of which 902 were upregulated (log2fold change > 1 and adjusted p-value < 0.05), and 511 were downregulated (log2fold change < –1 and adjusted p-value < 0.05) (Figure 2C). We identified 240 upregulated genes (Figure 2D), and 83 downregulated genes (Figure 2E) shared between the ivermectin and moxidectin-treated larvae.

To visualize the expression patterns of the differentially expressed transcripts in ivermectin– and moxidectin-treated larvae compared to controls, a heatmap of all expressed genes was generated and was sorted by gene expression (Figure 2F). We observed that approximately 11.22% of genes exhibited high expression levels, with normalized values between 10 to 15, indicated in yellow (10.94%), and between 15 to 25, indicated in red (0.28%). Conversely, a significant number of genes showed little to no expression, indicated in black (30%). The hierarchical clustering revealed two predominate clusters across different conditions, with further hierarchical sub-clustering within each.

To gain insight into the pathways and functions significantly associated with the observed changes in gene expression, a gene ontology (GO) enrichment analysis was performed on the differentially expressed genes (Figures 3-5). Differentially expressed genes were associated with GO terms categorized as biological processes (BP), molecular functions (MF), and cellular components (CC) and described in the following sections.

**Figure 3.**
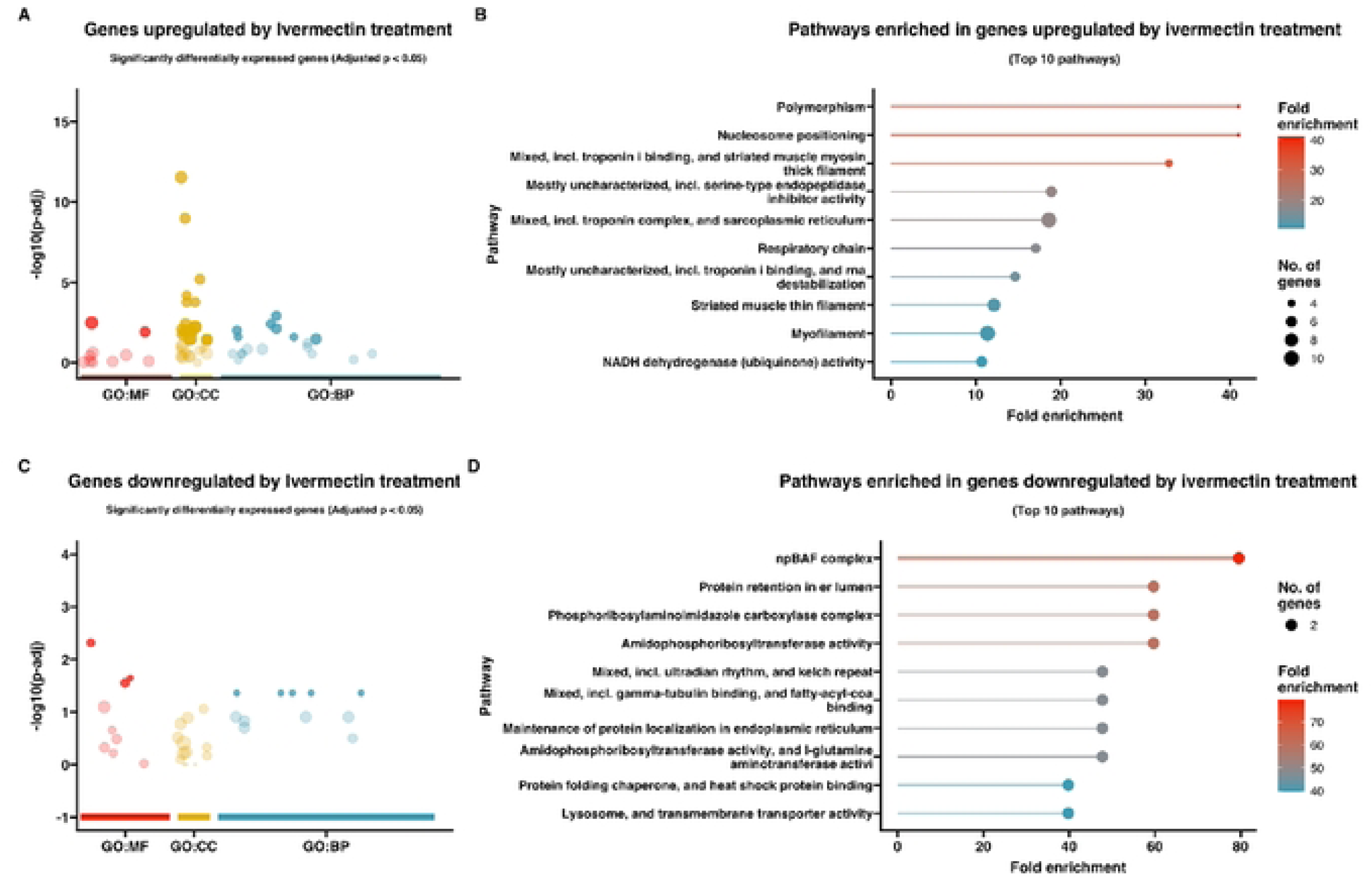
(**A&C**) Gene Ontology (GO) bubble plots of genes **(A)** upregulated and (C) do,vnregulated in *Toxocara canis* third-stage larvae (L3s) treated with ivermectin. The size of the bubble represents the number of genes associated with each GO term, and the color intensity of the bubble indicates the significance (adjusted p-value) of enrichment. GO terms are grouped by biological process (BP, blue), molecular function (MF, red), and cellular component (CC, yellow) categories; **(B&D)** Lollipop plots representing pathway enrichment analysis for the top IO most significantly enriched pathways for genes **(8)** upregulated and (C) downregulated in *T. canis* ivermectin-treated L3s. The length and color of the lollipops corresponds to fold enrichment, vith blue indicating lower fold enrichment and red indicating higher fold enrichment. The size of the dots represents the number of genes associate,dvith each pathway.

**Figure 4.**
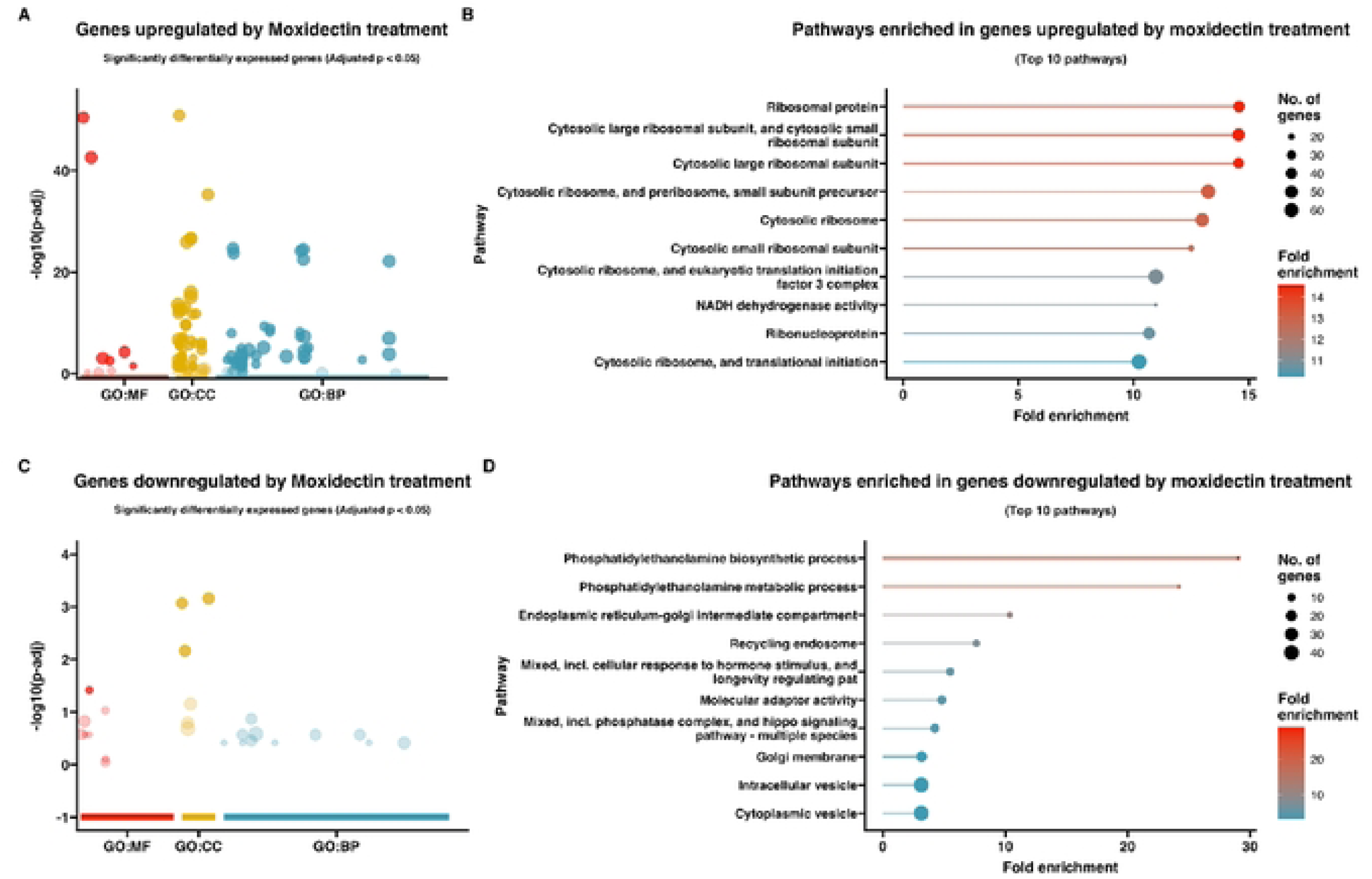
(**A&C**) Gene Ontology (GO) bubble plots of genes **(A)** upregulated and (C) do,vnregulated in *Toxocara canis* third-stage larvae (L3s) treated with moxidectin. The size of the bubble represents the number of genes associated with each GO term, and the color intensity of the bubble indicates the significance (adjusted p-value) of enrichment. GO terms are grouped by biological process (BP, blue), molecular function (MF, red), and cellular component (CC, yellow) categories; **(B&D)** Lollipop plots representing pathway enrichment analysis for the top IO most significantly enriched pathways for genes **(8)** upregulated and (C) downregulated in *T. canis* moxidectin-treated L3s. The length and color of the lollipops corresponds to fold enrichment, vith blue indicating lower fold enrichment and red indicating higher fold enrichment. The size of the dots represents the number of genes associated, vitheach pathway.

**Figure 5.**
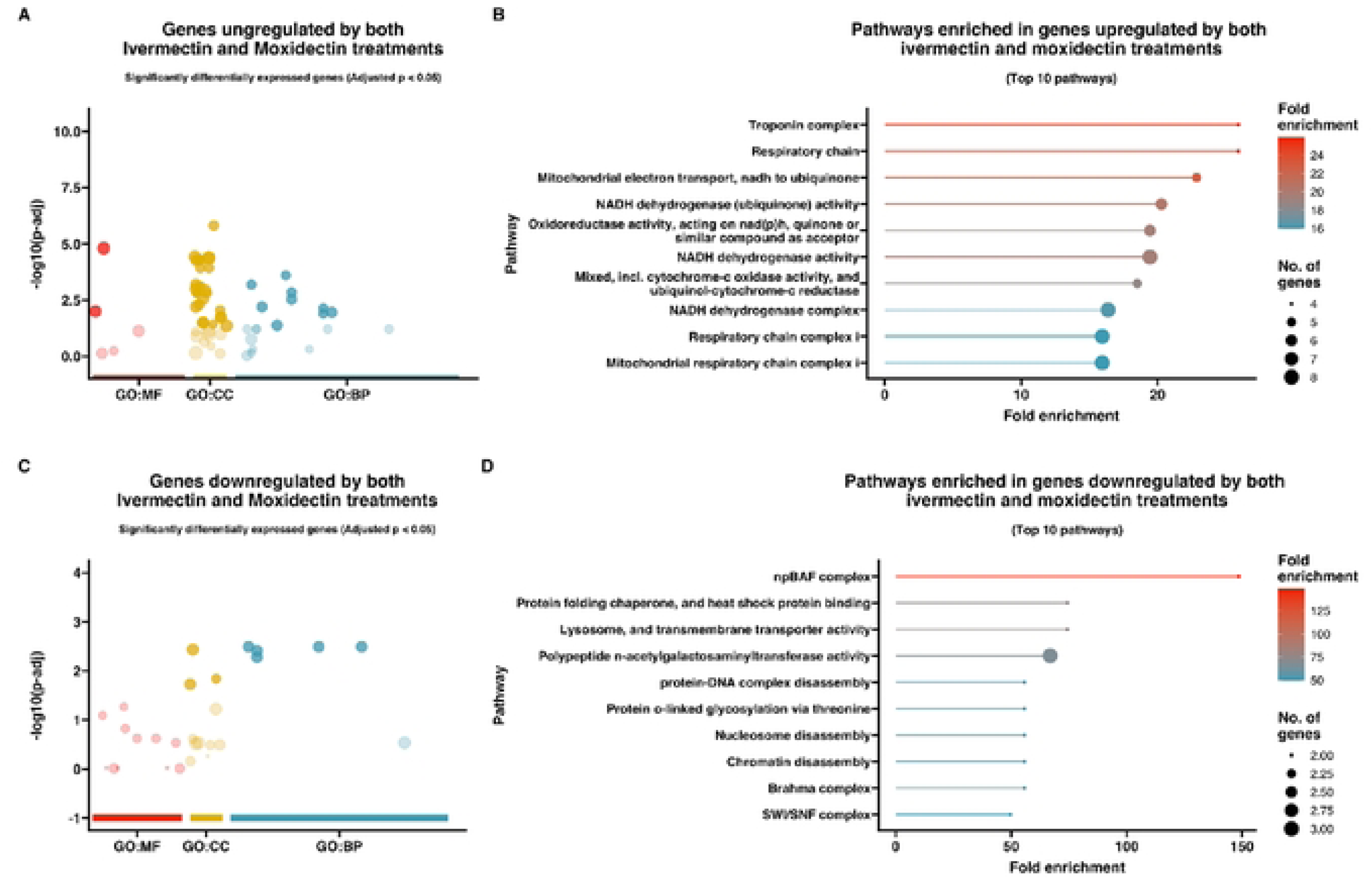
(**A&C**) Gene Ontology (GO) bubble plots of the overlapping **(A)** upregulated and (C) do,vnregulated shared genes between ivermectin– and moxidectin-treated *Toxocara canis* third-stage larvae (L3s). The size of the bubble represents the number of genes associated with each GO term, and the color intensity of the bubble indicates the significance (adjusted p-value) of enrichment. GO terms are grouped by biological process (BP, blue), molecular function (MF, red), and cellular component (CC, yello,v) categories; **(B&D)** Lollipop plots representing path,vay enrichment analysis for the top 10 most significantly enriched path,vays for the overlapping **(B)** upregulated and (C) do,vnregulated shared genes between ivermeetin– and moxidectin-trcatcd *1’. canis* L3s. The length and color of the lollipops corresponds to fold enrichment, with blue indicating lower fold enrichment and red indicating higher fold enrichment. The size of the dots represents the number of genes associated, vith each pathway.

### Response to ivermectin

The 453 *T. canis* genes upregulated in response to ivermectin exposure are listed in Supplemental Table 1A. Among the most upregulated genes in response to ivermectin exposure were a mediator of RNA polymerase II transcription subunit 13, *Tca-let-19* (Tcan_05892, log2fold change: 5, padj. 5.10E-03), glutamate dehydrogenase 2, mitochondrial, *Tca-GLUD2* (Tcan_03811, log2fold change: 3.9, padj. 2.32E-02), nuclear anchorage protein 1, *Tca-anc-1* (Tcan_00659, log2fold change: 3.5, padj. 1.09E-04), putative U5 small nuclear ribonucleoprotein helicase, *Tca-Y46G5.4* (Tcan_05430, log2fold change: 3.4, padj. 3.43E-03); additionally, there were six hypothetical proteins (Tcan_00335, Tcan_06163, Tcan_07262, Tcan_15031, Tcan_00594, and Tcan_00011), of which Tcan_06163 was identified as an ortholog of *C. elegans* T20G5.12.

Gene ontology enrichment analysis of the upregulated genes for larvae exposed to ivermectin (Supplemental Table 2A/ Figures 3A, 3B) revealed a significant enrichment of genes with GO terms associated with “polymorphism” (FDR = 6.6E-04), “nucleosome positioning” (FDR = 6.6E-04), “mixed, incl. troponin i binding, and striated muscle myosin thick filament” (FDR = 1.7E-04), “mostly uncharacterized, incl. serine-type endopeptidase inhibitor activity” (FDR = 4.3E-05), “mixed, incl. troponin complex, and sarcoplasmic reticulum” (FDR = 1.3E-08), “respiratory chain” (FDR = 3.7E-04), “mostly uncharacterized, incl. troponin i binding, and rna destabilization” (FDR = 46.6E-04), “striated muscle thin filament” (FDR = 42.7E-05) “myofilament” (FDR = 2.2E-06), and “NADH dehydrogenase (ubiquinone) activity” (FDR = 6.6E-04).

The 155 *T. canis* genes downregulated in response to ivermectin exposure are listed in Supplemental Table 1B. Among the most downregulated genes in response to ivermectin were a chondroitin proteoglycan 4, *Tca-cpg-4* (Tcan_10605, log2fold change: –4, padj. 3.76E-03), methylmalonyl-CoA epimerase, mitochondrial, *Tca-Mcee* (Tcan_12863, log2fold change: –3.3, padj. 2.18E-02), metalloproteinase inhibitor 2, *Tca-TIMP2* (Tcan_03888, log2fold change: –3.3, padj. 3.28E-02), putative oxidoreductase, *Tca-dhs-27* (Tcan_13635, log2fold change: –3.3, padj. 3.88E-02), amidophosphoribosyltransferase, *Tca-Prat* (Tcan_18524, log2fold change: –3.0, padj. 2.32E-03), nanos – like protein 2, *Tca-Nanos2* (Tcan_18885, log2fold change: –3.0, padj. 1.21E-02), and UDP-glucuronosyltransferase 2C1, *Tca-UGT2C1* (Tcan_05445, log2fold change: –2.9, padj. 6.19E-06). There were three hypothetical proteins (Tcan_15481, Tcan_14498, and Tcan_01557).

Gene ontology enrichment analysis of the downregulated genes for larvae exposed to ivermectin (Supplemental Table 2B/ Figures 3C, 3D) revealed a significant enrichment of genes with GO terms associated with “npBAF complex” (FDR = 2.1E-02), “amidophosphoribosyltransferase activity” (FDR = 2.8E-02), “protein retention in er lumen” (FDR = 2.8E-02), “phosphoribosylaminoimidazole carboxylase complex” (FDR = 2.8E-02)**, “**mixed, incl. ultradian rhythm, and kelch repeat” (FDR = 3.5E-02), “amidophosphoribosyltransferase activity, and l-glutamine aminotransferase activity” (FDR = 3.5E-02), “mixed, incl. gamma-tubulin binding, and fatty-acyl-coa binding” (FDR = 3.5E-02), “maintenance of protein localization in endoplasmic reticulum” (FDR = 3.5E-02), “lysosome, and transmembrane transporter activity” (FDR = 4.6E-02), and “protein folding chaperone, and heat shock protein binding” (FDR = 4.6E-02).

### Response to moxidectin

The 902 *T. canis* genes upregulated in response to moxidectin exposure are listed in Supplemental Table 3A. Among the most upregulated genes in response to moxidectin were a trypsin inhibitor, partial (Tcan_08038, log2fold change: 5.6, padj. 1.16E-04), glutamate dehydrogenase 2, mitochondrial, *Tca-GLUD2* (Tcan_03811, log2fold change: 4.9, padj. 1.12E-03), group XV phospholipase A2, *Tca-PLA2G15* (Tcan_15544, log2fold change: 4.7, padj. 7.56E-03), aspartyl protease inhibitor, partial, *Tca-API-3* (Tcan_02586, log2fold change: 4.3, padj. 9.96E-04), and protocadherin-16, *Tca-F32A5.4* (Tcan_07881, log2fold change: 3.8, padj. 3.56E-09). Five hypothetical proteins were also found (Tcan_01052, Tcan_00040, Tcan_15628, Tcan_00055, Tcan_00043), of which Tcan_00043 was identified as an ortholog to V-type proton ATPase subunit a in *Dirofilaria immitis* or *Bm33* in *Brugia malayi* [68, 69].

Gene ontology enrichment analysis of the upregulated genes for larvae exposed to moxidectin (Supplemental Table 4A/ Figure 4A, 4B) revealed a significant enrichment of genes with GO terms associated with “ribosomal protein” (FDR = 1.31E-38), “cytosolic large ribosomal subunit” (FDR = 9.45E-34), “cytosolic large ribosomal subunit, and cytosolic small ribosomal subunit” (FDR = 3.43E-45), “cytosolic ribosome, and preribosome, small subunit precursor” (FDR = 5.83E-53), “cytosolic ribosome” (FDR = 2.01E-48), “cytosolic small ribosomal subunit” (FDR = 1.45E-16), “cytosolic ribosome, and eukaryotic translation initiation factor 3 complex” (FDR = 3.65E-51), “NADH dehydrogenase activity” (FDR = 7.15E-13), “ribonucleoprotein” (FDR = 9.65E-34), and “cytosolic ribosome, and translational initiation” (FDR = 1.58E-49).

The 511 *T. canis* genes downregulated in response to moxidectin exposure are listed in Supplemental Table 3B. Among the most downregulated genes in response to moxidectin were an arginase, *Tca-rocF* (Tcan_06332, log2fold change: –4.2. padj. 2.42E-02), cuticlin-1, *Tca-cut-1* (Tcan_11029, log2fold change: –3.5, padj. 1.28E-02), alpha-aminoadipic semialdehyde synthase, mitochondrial, *Tca-AASS* (Tcan_18345, log2fold change: –3.4, padj. 1.32E-06), TRIO and F-actin-binding protein, partial, *Tca-Triobp* (Tcan_10839, log2fold change: –2.8, padj. 9.43E-03), lysine histidine transporter 1, *Tca-LHT1* (Tcan_17795, log2fold change: –2.8, padj. 1.20E-03), putative zinc finger protein F56D1.1, partial, *Tca-F56D1.1* (Tcan_16300, log2fold change: –2.6, padj. 3.47E-03), and methylmalonic aciduria and homocystinuria type D –like protein, mitochondrial, *Tca-Mmadhc* (Tcan_06175, log2fold change: –2.5, padj.1.43E-04). Three hypothetical proteins were identified (Tcan_04808, Tcan_02388, and Tcan_09119).

Gene ontology enrichment analysis of the downregulated genes for larvae exposed to moxidectin (Supplemental Table 4B/ Figure 4C, 4D) revealed a significant enrichment of genes with GO terms associated with “phosphatidylethanolamine biosynthetic process” (FDR = 0.00037204), “phosphatidylethanolamine metabolic process” (FDR = 0.000824851), “endoplasmic reticulum-golgi intermediate compartment” (FDR = 0.001363654), “recycling endosome” (FDR = 0.000785154), “mixed, incl. cellular response to hormone stimulus, and longevity regulating pat” (FDR = 0.001076538), “molecular adaptor activity” (FDR = 0.000374112), “mixed, incl. phosphatase complex, and hippo signaling pathway – multiple species” (FDR = 0.001056856), “golgi membrane” (FDR = 0.001403698), “cytoplasmic vesicle” (FDR = 6.59E-08), and “intracellular vesicle” (FDR = 6.59E-08).

### Shared analysis

To identify DEGs shared between ivermectin– and moxidectin-treated larvae, we performed a comparative analysis. DEGs shared in response to ivermectin and moxidectin exposure are listed in Supplemental Table 5. Among the top shared upregulated genes were glutamate dehydrogenase 2, mitochondrial, *Tca-GLUD2* (Tcan_03811, log2fold change: 4.4), protocadherin-16, *Tca-F32A5.4* (Tcan_07881, log2fold change: 3.0), cuticle collagen 40, *Tca-col-40* (Tcan_09436, log2fold change: 2.8), and two hypothetical proteins (Tcan_14010 and Tcan_15628). Tcan_14010 was identified as an ortholog to a “neo-calmodulin-like” protein in *Ylistrum balloti* (Ballot’s saucer scallop). Gene ontology enrichment analysis of upregulated shared genes (Supplemental Table 6A/ Figure 5A, 5B) revealed a significant enrichment of genes with GO terms associated with “troponin complex” (FDR = 3.7E-04), “respiratory chain” (FDR = 3.7E-04), “mitochondrial electron transport, nadh to ubiquinone” (FDR = 9.3E-05), “NADH dehydrogenase (ubiquinone) activity” (FDR = 2.7E-05), and “Oxidoreductase activity, acting on nad(p)h, quinone or similar compound as acceptor” (FDR = 3.5E-05).

Among the top shared downregulated genes were mitochondrial sodium/hydrogen exchanger 9B2, *Tca-SLC9B2* (Tcan_12528, log2fold change: –2.5), methylmalonic aciduria and homocystinuria type D –like protein, mitochondrial, *Tca-Mmadhc* (Tcan_06175, log2fold change: –2.3), T-cell activation inhibitor, mitochondrial, *Tca-TCAIM* (Tcan_14999, log2fold change: –2.3), as well as two hypothetical proteins (Tcan_00180 and Tcan_01557). Gene ontology enrichment analysis of downregulated shared genes (Supplemental Table 6B/ Figure 5C, 5D) revealed a significant enrichment of genes with GO terms associated with “npBAF complex” (FDR = 8.1E-03), “lysosome, and transmembrane transporter activity” (FDR = 1.8E-02), “protein folding chaperone, and heat shock protein binding” (FDR = 1.8E-02), “polypeptide n-acetylgalactosaminyltransferase activity” (FDR = 5.0E-03), and “nucleosome disassembly” (FDR = 2.3E-02).

### Genes associated with ML function and transport

To understand transcriptional changes in response to ML exposure in *T. canis* larvae, we identified 27 genes of interest annotated in the genome as ABCB transporters or GluCls and obtained their normalized expression (vst) values from the dataset. ABCB1 transporters of *T. canis* had been named previously [46] based on phylogenetic analysis following the recommendation of [70].

GluCls of *T. canis* have not been studied, and gene nomenclature has not been adequately addressed. To clarify gene naming for *T. canis* GluCls, we identified seven GluCls in the transcriptome of *T. canis* and assigned gene names to them (Table 2) using a maximum likelihood phylogenetic tree (Figure 6). The phylogenetic analysis showed that orthologs of GluCls in *T. canis* clustered closely with those of other described nematodes, forming well-supported clades. Additionally, we identified one *T. canis* gene (GenBank Accession KHN83224) that was annotated as a GluCl; however, phylogenetic analysis revealed that it clustered within a distinct clade alongside other nematode ligand-gated ion channels. Thus, we included 19 ABCB transporter genes, seven GluCls, and one ligand-gated ion channel in our analysis.

**Figure 6.**
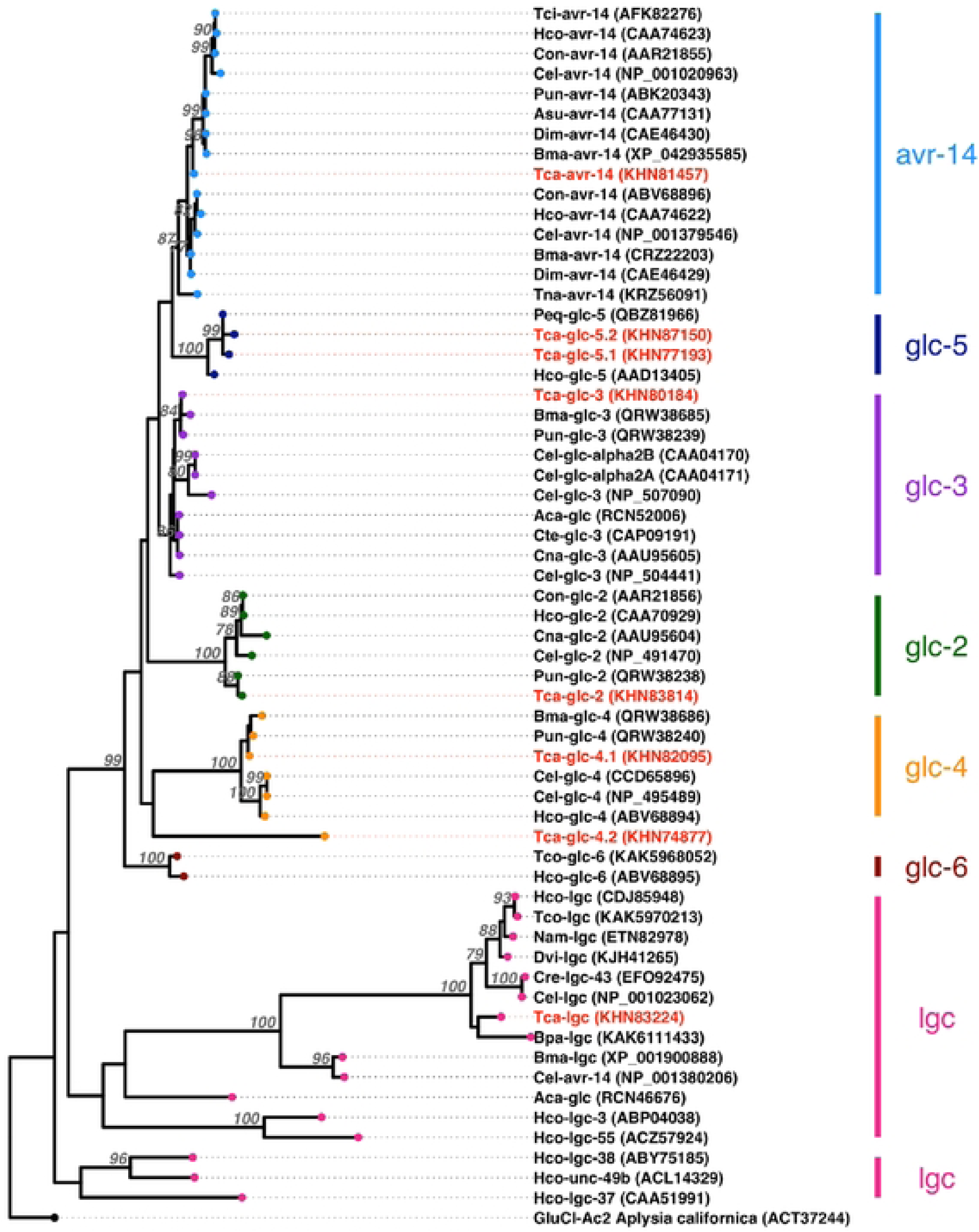
A maximum-likelihood phylogenetic tree of *Toxocara canis* glutamate-gated chloride channels (GluCls) and related nematode sequences obtained from GenBank. The tree was constructed using PhyML, vith the SMS to dctenninc the best fitting model and visualized using iTOL and ggtrce. Branch support values were calculated from 1000 bootstrap replicates, with values> 75 shown. The tree is rooted using a GluCl sequence from *Aplysia californica* (California sea hare). The tree revealed six distinct GluCl clades (avr-14, glc-5, glc-3, glc-2, glc-4, and glc-6), as, veil as distinct clades for nematode ligand­ gated channels (lgc).

**Table 2.** Nomenclature for GluCl protein sequences in *Toxocara canis*.

| GenBank accession number<br>(Protein database) | Assigned gene names |
| --- | --- |
| KHN83814 | *Tca-glc-2* |
| KHN87150 | *Tca-glc-5.2* |
| KHN83224 | *Tca-lgc* |
| KHN82095 | *Tca-glc-4.1* |
| KHN81457 | *Tca-avr-14* |
| KHN80184 | *Tca-glc-3* |
| KHN77193 | *Tcaglc-5.1* |
| KHN74877 | *Tca-glc-4.2* |

**Table 3.** Nomenclature for ABCB protein sequences in *Toxocara canis*.

| GenBank accession number (Protein database) | Assigned gene names | Full or partial | Reference |
| --- | --- | --- | --- |
| KHN80157 | *Tca-Pgp-2* | Full | [46] |
| KHN78383 | *Tca-Pgp-3* | Partial | [46] |
| KHN77280 | *Tca-Pgp-9.1* | Partial | [46] |
| KHN76604 | *Tca-Pgp-9.2* | Partial | [46] |
| KHN84692 | *Tca-Pgp-10* | Full | [46] |
| KHN73709 | *Tca-Pgp-11.1* | Full | [46] |
| KHN87227 | *Tca-Pgp-11.2* | Full | [46] |
| KHN89031 | *Tca-Pgp-11.3* | Full | [46] |
| KHN87070 | *Tca-Pgp-13.1* | Full | [46] |
| KHN87068 | *Tca-Pgp-13.2* | Full | [46] |
| KHN80315 | *Tca-Pgp-16.1* | Full | [46] |
| KHN80341 | *Tca-Pgp-16.2* | Full | [46] |
| KHN86334 | *Tca-Pgp-16.3/3.2* | Full | [46] |
| KHN88383 | *Tca_abcb8* | Partial |  |
| KHN84272 | *Tca-abcb1* | Partial |  |
| KHN83466 | *Tca-abcb7* | Partial |  |
| KHN73719 | *Tca-abcb9* | Partial |  |
| KHN73722 | *Tca-abcb9* | Partial |  |
| KHN76663 | *Tca-abcb9* | Partial |  |

We analyzed the expression of the identified GluCls and ABCBs [46] (Figure 7A). In our initial DE analysis of all 18,596 genes, GluCls were not found among the DEGs and only ABCB9 was upregulated in the moxidectin-treated larvae compared to controls. Upon further analysis using Shapiro-Wilk and Tukey multiple comparisons tests, we observed differences in the expression of seven out of the 27 genes of interest (Figure 7B). The Shapiro-Wilk test identified 25 out of the 27 genes were normally distributed (p > 0.05). Subsequently, Tukey’s test identified that *Tca-glc-3* was significantly downregulated in ivermectin-treated larvae compared to controls (p = 0.036). Of the ABCBs, compared to controls, *Tca-abcb1*, *Tca-abcb7*, *Tca-Pgp-11.2*, and *Tca-Pgp-13.2* were downregulated in moxidectin-treated larvae (p = 0.042, 0.024, 0.0448, and 0.0071, respectively). *Tca-Pgp-11.2* and *Tca-Pgp-13.2* were also downregulated in ivermectin-treated larvae compared to controls (p = 0.043 and 0.0071, respectively). Conversely, *Tca-abcb9.1* was upregulated in moxidectin-treated larvae compared to controls (p = 0.0076). Finally, *Tca-Pgp-11.3* was upregulated in moxidectin-treated larvae compared to both the controls and ivermectin-treated larvae (p = 0.0092 and 0.0488, respectively).

**Figure 7.**
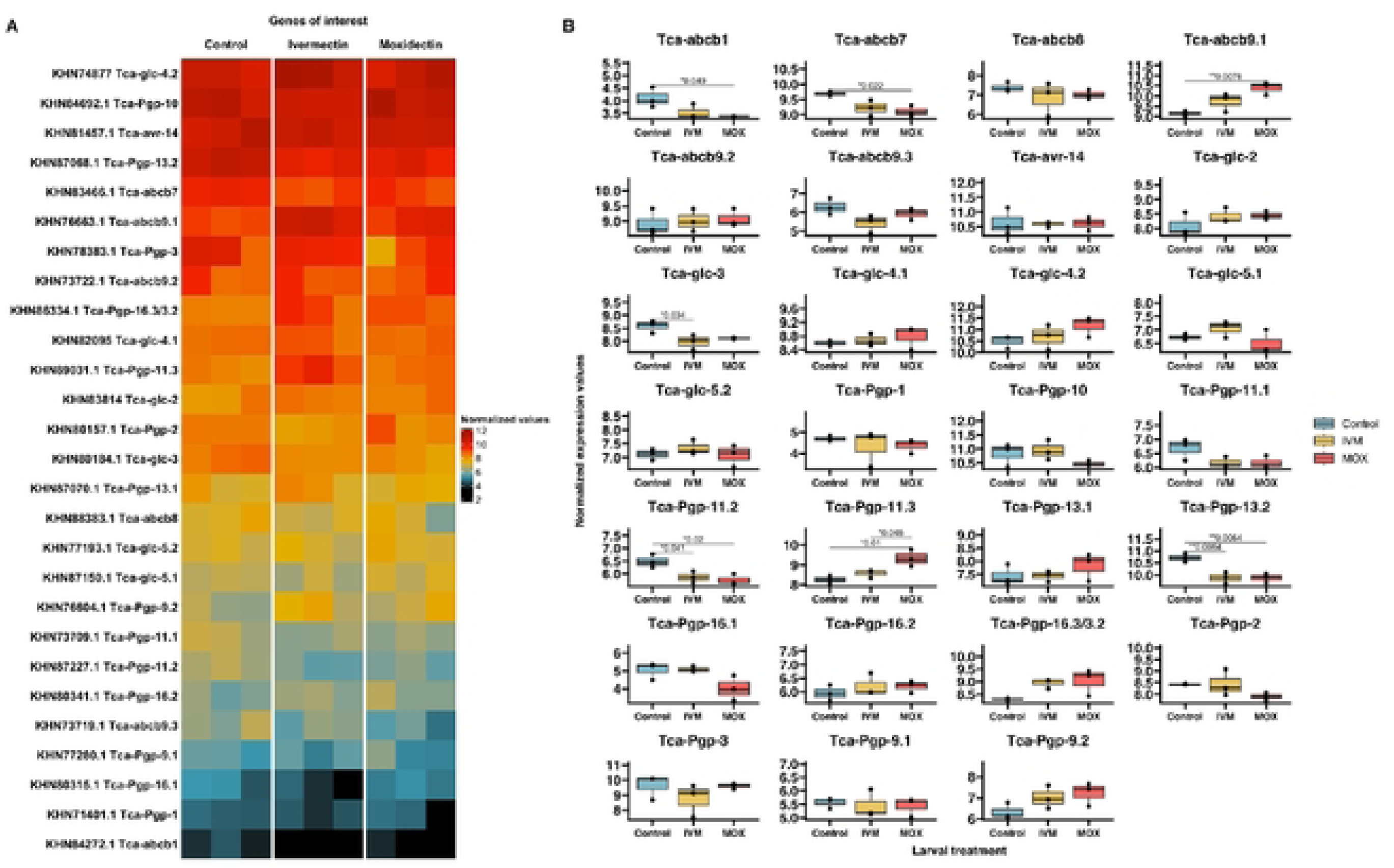
(**A**) Heatmap of normalized expression (vst) values for all 27 *Toxocara canis* genes of interest, including six glutamate-gated chloride channels (GluCls}, 19 ATP-Binding Cassette Transporters, subfamily B (13 of which arc P-glycoproteins (Pgps)}, and one ligand-gated channel (lgc), for control, ivermectin-, and moxidectin-treated third-stage larvae (L3s). Color intensity represents normalized expression values, with the scale ranging from black (lowest expression) to red (highest expression). **(B)** Boxplots of normalized expression (vst) values for the 27 T. canis genes of interest. The y-axis represents normalized expression. Statistical significance between treatment groups was assessed using Tukey’s multiple comparison. Control L3s arc represented as blue, ivcrmectin-treated L3s as yellow, and moxidectin-treated L3s as red shaded boxes.

## Discussion

*T. canis* infections remain a persistent threat to both humans and animals globally, despite the availability of effective pharmaceuticals for treating adult infections. Somatic larvae within canine hosts serve as the primary reservoir for propagating the lifecycle and are not cleared using MLs at approved doses [27]. The mechanisms behind the tolerance of *T. canis* larvae to MLs are still poorly understood. In this study, we assessed the effects of the MLs on *T. canis* third-stage larvae *in vitro* using transcriptomics, focusing on DEGs and the expression levels of genes associated with ML action and resistance in nematodes, specifically GluCls and P-gps.

To increase the rigor of the study, we used three replicates of larvae hatched from eggs from different adult worms and subjected them to 10 μM ivermectin, moxidectin, or no drugs (RPMI-1640) for 12 hours and performed RNA-seq on the Illumina platform. We observed a high intragroup Pearson correlation of mRNA transcripts in the control group (0.94–0.99) indicating strong homogeneity among the samples. The ivermectin group showed greater variability (0.71–0.95), while the moxidectin group (0.97–0.98) demonstrated a consistency similar to the control group. Furthermore, we observed significant differential expression of genes in larvae exposed to ivermectin (609 genes) and moxidectin (1,413 genes) compared to controls. Of these, 453 were upregulated and 155 were downregulated in response to ivermectin exposure, while 902 were upregulated and 511 were downregulated in response to moxidectin exposure. Among these, 240 were shared between the two MLs in the upregulated genes and 83 in the downregulated gene set.

The consensus site of action of the MLs are the GluCls. GluCls are pentameric channels composed of heterogeneous or homogeneous subunits. MLs bind to these channels between the M1 and M3 transmembrane domains, specifically at the Leu217 and Ile222 sites on M1 [71]. GluCls have been extensively studied in *Ascaris suum* [72], *C. elegans* [71, 73–78], *H. contortus* [79–84] and *Cooperia oncophora* [85]. However, GluCls have not been well characterized in *T. canis*. The *T. canis* genome encodes seven GluCls and these were expressed in the larval stages in this study. We clarified the nomenclature of GluCls using phylogenetic analysis (Figure 6, Table 2) as recommended by [70]. While GluCls were not among the DEGs in our initial analysis, upon further exploration we found that *Tca-glc-3* was downregulated in larvae exposed to ivermectin (Figure 7B). Similar transcriptomic analysis in ivermectin-resistant strains of *H. contortus* from China showed that *Hco-glc-5* was downregulated compared to susceptible strains from Australia [32]. However, studies have shown that *Hco-glc-5* has no evidence of linkage to an ivermectin resistance phenotype in United Kingdom *H. contortus* isolates [86, 87]. Further research is needed to clarify the specific roles of GluCls in ML resistance.

ML resistance and tolerance in parasitic nematodes have been associated with P-gps (Synonym: ABCB1) [88–91]. P-gps have two ATP binding domains and two transmembrane domains [92]. Since P-gps are monocistronic, each fully annotated gene corresponds to a single protein. Other ABCB transporters include subclasses A, C, D, E, F, G, and H. The *T. canis* genome encodes thirteen P-gps and six other ABCB genes. We assessed the expression level of the nineteen genes, including the thirteen named P-gp genes [46] and six ABCB genes annotated in the genome. While P-gps (ABCB1s) were not among the DEGs, ABCB9 was upregulated in the moxidectin treatment group. Upon further analysis, differences in the expression of a further five ABCB transporter genes were observed. In response to MLs, *Tca-Pgp-11.2* and *Tca-Pgp-13.2*, were downregulated, while *Tca-Pgp-11.3* was upregulated. This is in agreement with previous findings that *T. canis* larvae exposed to ivermectin downregulate the expression of *Tca-Pgp-13.1* [46].

Beyond GluCl and P-gp genes of interest, additional pathways impacted by MLs include those involved in the regulation of transcription, energy production, neuronal activity, essential physiological functions like reproduction, expression of genes responsible for excretory/secretory (ES) molecules, responses to host signals, and mechanisms for detoxification.

MLs influence gene transcription by modulating mechanisms that control transcriptional regulation. In this study, ivermectin was observed to upregulate a mediator of RNA polymerase II transcription subunit 13, *Tca-let-19* (Tcan_05892). This gene is part of the mediator complex that governs gene expression by influencing transcriptional initiation and regulation [93]. Similar findings in osteosarcoma models suggest that this mediator complex plays a critical role in promoting tumor growth and cellular stress responses [94]. In addition, ivermectin upregulated the putative U5 small nuclear ribonucleoprotein helicase, *Tca-Y46G5.4* (Tcan_05430), which is involved in pre-mRNA splicing and post-transcriptional RNA modification. Moxidectin, on the other hand, downregulated the putative zinc finger protein F56D1.1 (Tcan_16300), a protein known to bind DNA, RNA, and proteins, playing a vital role in gene regulation, DNA repair, and other essential cellular processes. GO enrichment analysis provided further evidence that several transcription-related processes were significantly affected, including the upregulation of pathways associated with nucleosome positioning, RNA destabilization, ribosomal activity, and translational initiation in larvae exposed to MLs (Figures 3B & 4B). These demonstrate the impact MLs have either directly or indirectly on transcriptional regulation.

ML exposure impacted energy metabolism by modulating the expression of key metabolic enzymes. Ivermectin and moxidectin upregulated mitochondrial glutamate dehydrogenase 2 (*Tca-GLUD2*, Tcan_03811), an enzyme studied in the parasitic nematodes *Teladorsagia circumcincta* and *Haemonchous contortus* that is crucial for nitrogen and energy metabolism, converting glutamate to α-ketoglutarate and ammonia [95, 96]. In contrast, ivermectin downregulated methylmalonyl-CoA epimerase, mitochondrial (*Tca-Mcee*, Tcan_12863), an enzyme involved in the metabolism of fatty acids and amino acids in *C. elegans* [97]. Ivermectin also downregulated amidophosphoribosyltransferase (*Tca-Prat*, Tcan_18524), which catalyzes the first step in purine biosynthesis and has been shown to be differentially expressed in *Wolbachia* during development of *Brugia malayi* male and female adults [98]. Moxidectin downregulated other enzymes involved in amino acid and vitamin metabolism, including a lysine histidine transporter 1 (*Tca-LHT1*, Tcan_17795), a membrane protein responsible for the transport of lysine and histidine; a mitochondrial alpha-aminoadipic semialdehyde synthase, (*Tca-AASS*, Tcan_18345), shown to be involved in lysine degradation in *C. elegans* [99]; and a mitochondrial methylmalonic aciduria and homocystinuria type D-like protein, (*Tca-Mmadhc*, Tcan_06175), which plays a role in Vitamin B12 metabolism. GO enrichment analysis further revealed that DEGs in response to MLs were related to metabolic pathways, including upregulation of those associated with NADH dehydrogenase activity and the respiratory chain (Figures 3B & 4B), indicating MLs may affect *T. canis’* ability to metabolize energy efficiently.

MLs play a role in neuronal structure and function by modulating the transcription of key genes. Ivermectin upregulated nuclear anchorage protein 1 (*Tca-anc-1*, Tcan_00659), which is essential for maintaining nucleus-cytoskeleton interactions and stabilizing neuron positions in *C. elegans* [100]. Moxidectin similarly upregulated protocadherin-16 (*Tca-F32A5.4*, Tcan_07881), a protein likely involved in cell-cell adhesion and neuronal development, contributing to tissue morphogenesis and homeostasis. In contrast, moxidectin downregulated cuticlin-1 (*Tca-cut-1*, Tcan_11029), described in the parasitic nematode *Ascaris suum* as a key structural protein of the epicuticle that is rich in proline and alanine [101, 102]. In the parasitic nematode *B. malayi*, downregulation of cuticle collagen genes in response to ivermectin has been studied [103]. Furthermore, in ivermectin-resistant *H. contortus*, studies have shown that cuticle collagen genes are downregulated when compared to susceptible isolates [104]. Additionally, moxidectin downregulated TRIO and F-actin-binding protein (*Tca-Triobp*, Tcan_10839). In *C. elegans*, UNC-73, a TRIO homolog, is involved in regulating cytoskeletal and neuronal development [105]. Finally, the differential expression of genes corresponding to the GO enrichment pathways for the troponin complex, sarcoplasmic reticulum, and myofilament formation (Figure 3B) suggest MLs may disrupt normal neuronal processes.

MLs impact the transcription of genes involved in physiological functions, including reproduction. Ivermectin downregulated chondroitin proteoglycan 4 (*Tca-cpg-4*, Tcan_10605), which may play a role in reproductive processes, similar to chondroitin proteoglycan 2 (*Tca-cpg-2*); *Tca-cpg-2* in adult female *T. canis* is crucial for oogenesis and embryogenesis and is enriched in germline tissues [106]. Additionally, ivermectin downregulated nanos-like protein 2 (*Tca-Nanos2*, Tcan_18885), which is involved in germ cell development. GO enrichment analysis showed that MLs downregulated pathways involved in cellular responses to hormone stimuli and longevity-regulating processes which could suggest that MLs interfere with essential biological functions (Figure 4B).

MLs impact the transcription of genes that encode excretory/secretory (ES) and other molecules crucial for host-parasite interactions. Ivermectin downregulated a metalloproteinase inhibitor 2 (*Tca-TIMP2*, Tcan_03888). In *Ancylostoma caninum*, the metalloproteinase inhibitor AcTMP is speculated to regulate host matrix metalloproteinases or host neutrophil collagenase activity to influence tissue repair and inflammation at the intestinal mucosa attachment site [107]. Moxidectin, on the other hand, upregulated multiple protective factors, including a trypsin inhibitor (Tcan_08038) and an aspartyl protease inhibitor (*Tca-API-3*, Tcan_02586), which may protect against host-specific proteolytic enzymes and play a role in the host immune response and infection, as seen in *A. suum* [108], *H. contortus* [109], and *Ostertagia ostertagi* [110]. Additionally, moxidectin upregulates a group XV phospholipase A2 (*Tca-PLA2G15*, Tcan_15544). In parasitic nematodes, phospholipase A2 (PLA2) plays crucial roles in pathogenesis by contributing to host tissue penetration, modulating host immune responses, and facilitating nutrient acquisition [111]. These enzymes disrupt host cell membranes, release signaling molecules that manipulate host immune pathways, and help the parasites maintain cellular integrity and repair, thereby enhancing their survival and ability to infect the host [111]. Finally, moxidectin downregulated arginase (*Tca-rocF*, Tcan_06332), which normally competes with nitric oxide synthase for L-arginine, reducing nitric oxide production and aiding immune evasion [82]. The differential expression of genes responsible for the production of ES molecules may indicate that MLs alter the ability of *T. canis* to interact within the host.

MLs influence *T. canis*’ ability to respond to host cues. Ivermectin downregulated a putative oxidoreductase, *Tca-dhs-27* (Tcan_13635), an enzyme involved in redox reactions that is crucial for maintaining cellular redox homeostasis. Interestingly, a similar oxidoreductase is upregulated in *Anisakis simplex* larvae in response to ivermectin [112]. This difference between species warrants further investigation into the role of MLs in parasitic responses to host cues.

MLs affect the transcription of genes involved in *T. canis’* elimination mechanisms, altering detoxification pathways. Ivermectin downregulated UDP-glucuronosyltransferase 2C1, *Tca-UGT2C1* (Tcan_05445), an enzyme for detoxifying both endogenous and exogenous compounds. UDP-glucuronosyltransferases facilitate the glucuronidation process, which makes drugs and xenobiotics more water-soluble, enhancing their excretion [113]. The downregulation of a detoxification enzyme is surprising and suggests MLs may impair *T. canis*’ ability to effectively detoxify harmful compounds.

MLs affect the transcription of several hypothetical genes, which have unknown functions (distinct from pseudogenes). Ivermectin upregulated Tcan_06163, identified as an ortholog of *C. elegans* T20G5.12, though its function remains unknown. Additionally, ivermectin upregulated Tcan_00335, a SET domain family protein. In *C. elegans,* SET domain proteins play a crucial role in regulating gene expression through histone methylation, influencing key processes such as development, reproduction, chromatin remodeling, and lifespan control [114–116]. Ivermectin downregulated Tcan_14498, a domain-containing protein. In *C. elegans*, this domain has no known functions (pfam PF01682) but has been associated with immunoglobulin and fibronectin domains, and lipases [117]. Moxidectin upregulated Tcan_00043, which was identified as either an ortholog of the V-type proton ATPase subunit A in *D. immitis* or Bm33 in *B. malayi*. V-type ATPases are involved in acidifying intracellular endosomes, lysosomes, and Golgi-derived vesicles and hyperpolarizing membranes through ATP hydrolysis to produce a proton gradient[118]. Bm33 is an aspartyl protease inhibitor thought to play a role in immune evasion by inhibiting host proteases[119]. Moxidectin also upregulated Tcan_00055, identified as an ortholog to a regulator of rDNA transcription protein 15 in *Trichinella papuae* [120]. Hypothetical proteins without identifiable orthologs were also observed. The lack of functional annotations for these genes limits our understanding of their precise roles within *T. canis* and therefore limits our ability to predict the impact of MLs.

Some limitations of this study include the use of a small sample size and only a 12-hour drug exposure period, which may not fully capture the range of biological responses to ML treatments.

Additionally, the study was conducted *in vitro* using a single 10 µM dose which limits the generalizability of the findings to real-world scenarios where drug exposure may vary. Another limitation is the absence of a well-annotated *Toxocara canis* chromosome-level genome assembly, making accurate mapping more challenging. Future research should investigate the effects of multiple doses over extended periods, explore alternative drugs, and incorporate multi-omics approaches, such as proteomics, to provide a more comprehensive understanding of the biological mechanisms involved. Additionally, mouse models could be employed to study and compare transcriptomic results from untreated and ML-treated larvae *in vivo* to *in vitro* studies, facilitating the interpretation of the impact of MLs on somatic L3s within a host.

### Concluding statement

In conclusion, this study provides valuable insights into how MLs like ivermectin and moxidectin impact the transcription of various genes associated with essential biological processes in *T. canis* hatched L3s. In L3s treated *in vitro* with ivermectin or moxidectin, we identified 608 and 1,413 DEGs, respectively, of the 18,596 genes in the genome. Among the transcriptome were genes known to be associated with ML resistance, including GluCls and P-gps, which remain of significant interest as potential targets for therapeutic interventions. We also identified DEGs of potential interest that could confer ML tolerance in *T. canis* L3s, which could serve as valuable candidates for future research and exploration. Future studies should employ the use of multi-omics approaches to capture the full spectrum of molecular changes induced by MLs. Additionally, investigating the effects of MLs at multiple doses over extended periods will help determine long-term impacts on *T. canis* larvae. Researchers should also consider exploring the impacts of alternative anthelmintic compounds to identify treatments that may overcome tolerance mechanisms associated with MLs. Lastly, comparing *in vivo* larvae to *in vitro* larvae studies will help clarify the specific gene expression changes and physiological effects induced by MLs within a host environment. Together, these approaches will provide a comprehensive understanding of MLs’ effects on *T. canis*.

## Ethics statement

All protocols were approved by the Institutional Biosafety Committee at Kansas State University (IBC-1516-VCS)

## Data Availability

The dataset generated and analyzed during this study is available through the National Center for Biotechnology Information (NCBI; www.ncbi.nlm.nih.gov) under accession numbers SAMN38303355 – SAMN38303369 (Bioproject: PRJNA1041894).

## Acknowledgements

We acknowledge the assistance of the Kansas State Veterinary Diagnostic Laboratory necropsy section and Laura Stein and her staff with the Dodge City Animal Control for providing adult *T. canis*. We would like to acknowledge Drs. Patricia Assato and Jayme Souza-Neto of the Center for Emerging and Zoonotic Infectious Diseases (CEZID) Molecular and Cellular Biology Core for their support with sequencing. We acknowledge the public servers at usegalaxy.org which is supported by NIH and NSF Grants HG006620, 1661497, and 1929694.

## Author contributions

Conceptualization: T.A.Q., M.T.B. and J.R.J.; Data curation: T.A.Q. and J.R.J.; Formal analysis: T.A.Q. and J.R.J.; Funding acquisition: J.R.J.; Investigation: T.A.Q. and J.R.J.; Methodology: T.A.Q. and J.R.J.; Project administration: J.R.J.; Software: T.A.Q. and J.R.J.; Supervision: J.R.J.; Validation: T.A.Q. and J.R.J.; Visualization: T.A.Q. and J.R.J.; Writing – original draft: T.A.Q. and J.R.J.; Writing – review & editing: T.A.Q., M.T.B. and J.R.J.

